# Environmental conditions and diffusion-limited microbial transfer drive specific microbial communities detected at different sections in oil-production reservoir with water-flooding

**DOI:** 10.1101/2020.12.07.415646

**Authors:** Peike Gao, Huimei Tian, Guoqiang Li, Feng Zhao, Wenjie Xia, Ji-Dong Gu, Jianjun Le, Ting Ma

**Affiliations:** College of Life Sciences, Qufu Normal University, Qufu, Shandong, 273165, The People’s Republic of China; College of Forestry, Shandong Agricultural University, Tai’an, Shandong, 271018, The People’s Republic of China; College of Life Sciences, Nankai University, Tianjin, 300071, The People’s Republic of China; Environmental Engineering, Guangdong Technion - Israel Institute of Technology, 241 Daxue Road, Shantou, Guangdong, 515063, The People’s Republic of China; Exploration and Development Research Institute, PetroChina Daqing Oilfield Limited Company, Daqing, Heilongjiang, 163000, The People’s Republic of China

**Keywords:** Environmental selection, Metabolic profiles, Oilfield production facilities, Petroleum reservoir

## Abstract

This study investigated the distribution of microbial communities in the oilfield production facilities of a water-flooding petroleum reservoir and the roles of environmental variation, microorganisms in injected water, and diffusion-limited microbial transfer in structuring the microbial communities. Similar bacterial communities were observed in surface water-injection facilities dominated by aerobic or facultative anaerobic Betaproteobacteria, Alphaproteobacteria, and Flavobacteria. Distinct bacterial communities were observed in downhole of the water-injection wells dominated by Clostridia, Deltaproteobacteria, Anaerolineae, and Synergistia, and in the oil-production wells dominated by Gammaproteobacteria, Betaproteobacteria, and Epsilonproteobacteria. *Methanosaeta, Methanobacterium*, and *Methanolinea* were dominant archaeal taxa in the water-injection facilities, while the oil-production wells were predominated by *Methanosaeta*, *Methanomethylovorans*, and *Methanocalculus*. Energy, nucleotide, translation, and glycan biosynthesis metabolisms were more active in the downhole of the water-injection wells, while bacterial chemotaxis, biofilm formation, two-component system, and xenobiotic biodegradation was associated with the oil-production wells. The number of shared OTUs and its positive correlation with formation permeability revealed differential diffusion-limited microbial transfer in oil-production facilities. The overall results indicate that environmental variation and microorganisms in injected water are the determinants that structure microbial communities in water-injection facilities, and the determinants in oil-bearing strata are environmental variation and diffusion-limited microbial transfer.

**IMPORTANCE:** Water-flooding continually inoculates petroleum reservoirs with exogenous microorganisms, nutrients, and oxygen. However, how this process influences the subsurface microbial community of the whole production process remains unclear. In this study, we investigated the spatial distribution of microbial communities in the oilfield production facilities of a water-flooding petroleum reservoir, and comprehensively illustrate the roles of environmental variation, microorganisms in injected water, and diffusion-limited microbial transfer in structuring the microbial communities. The results advance fundamental understanding on petroleum reservoir ecosystems that subjected to anthropogenic perturbations during oil production processes.

## INTRODUCTION

Petroleum reservoirs contain microorganisms with diverse phylogenetic affiliations and metabolic characteristics, including hydrocarbon-degrading bacteria, fermentative bacteria, sulfate-reducing bacteria, iron-reducing bacteria, acetogens, and methanogens, among others (1, 2). The main microbial processes prevailing in petroleum reservoir ecosystems include hydrocarbon degradation, nitrate reduction, sulfate reduction, fermentation, acetogenesis, methanogenesis, and iron and manganese reduction (3, 4). With the increasing global demand for crude oil, research in the field of petroleum reservoir microbial communities has attracted increasing attention due to its great potential in improving oil production processes, such as microbiologically enhanced oil recovery (5–11) and microbiologically-prevented reservoir souring and equipment corrosion (12–16).

In recent decades, a large number of studies have investigated the composition of microbial communities in global petroleum reservoirs (17–21). Given the pronounced differences in inherent conditions among the reservoirs, in particular key microbial growth limiting factors, such as temperature (13, 22), salinity (23, 24), and pH (25), a variety of microbial ecological patterns with high variability in community composition have been identified in these ecosystems. Furthermore, studies are increasingly indicating that microbiologically-improved oil production processes are closely related to changes in the microbial communities of reservoirs, such as altered microbial abundance and composition (5, 26–29). This is of great significance to the study of the microbial communities inhabiting petroleum reservoirs.

Petroleum reservoir ecosystems are subject to extreme anthropogenic perturbations during oil exploration and oil production processes, such as drilling, workover, and the application of secondary and tertiary oil recovery techniques, all of which introduce new electron acceptors, donors, and exogenous microbes into reservoir environments. Recently, Vigneron et al. (2017) elucidated the succession of microbial communities that occur over the production lifetime of an offshore petroleum reservoir (19). Their results expanded our current knowledge regarding the shifts of reservoir microbial communities in different stages of oil exploitation. However, little is known about the microbial communities in oilfield production facilities. Research in this field has profound consequences on our understanding of the microbial community distribution in reservoirs and the factors driving changes, and will improve our ability to predict and regulate reservoir microbial communities to microbiologically improve oil production processes.

Petroleum and formation water are usually pushed to the surface by pressure naturally found within a reservoir, known as primary recovery. Subsequently, water injection is an efficient and inexpensive secondary recovery process that is widely used to maintain reservoir pressure and achieve a higher oil-production level. The injected water generally consists of recycled water produced from oil-production wells and make-up water consisting of seawater, river water, or underground water, which contains large amounts of inorganic ions (e.g. nitrate and sulfate), dissolved oxygen, and a mass of exogenous microorganisms (17, 19, 22, 30). The water-flooding process results in the continual inoculation of reservoirs with exogenous microorganisms, nutrients, and oxygen, which will likely alter reservoir geochemistry either temporarily or permanently, and significantly influence the subsurface microbial community.

In water-flooding oil reservoirs, the injected water from water injection stations flows into water-injection wells, then into oil-bearing strata, and outflows from oil-production wells with crude oil (Fig. 1). Before water flows into oil-bearing strata, the microorganisms in the injected water may be easily transferred from the water-injection station to the wellheads and downhole of the water-injection wells. It is not hard to speculate that the exogenous microorganisms in the injected water and environmental variation within habitats rather than diffusion-limited microbial transfer may play a more crucial role in structuring the microbial communities in the pipelines. Once the injected water flows from the downhole of water-injection wells into the oil-bearing strata and then into the oil-production wells, environmental variation within habitats and diffusion-limited microbial transfer may lead to microbial community assembly processes. Several studies have observed considerable uniformity among the microbial communities inhabiting geographic-adjacent or -isolated oil-production wells (17, 18, 31, 32). However, whether the exogenous microorganisms are able to enter oil-bearing strata and reach oil-production wells remains a subject of debate, let alone their effects on subsurface microbial communities. Therefore, a systematic study of the microbial communities is needed to elucidate the roles of environmental variables within habitats, including the microorganisms in injected water and diffusion-limited microbial transfer, in the structuring of the microbial communities in petroleum reservoirs.

**Fig. 1.**
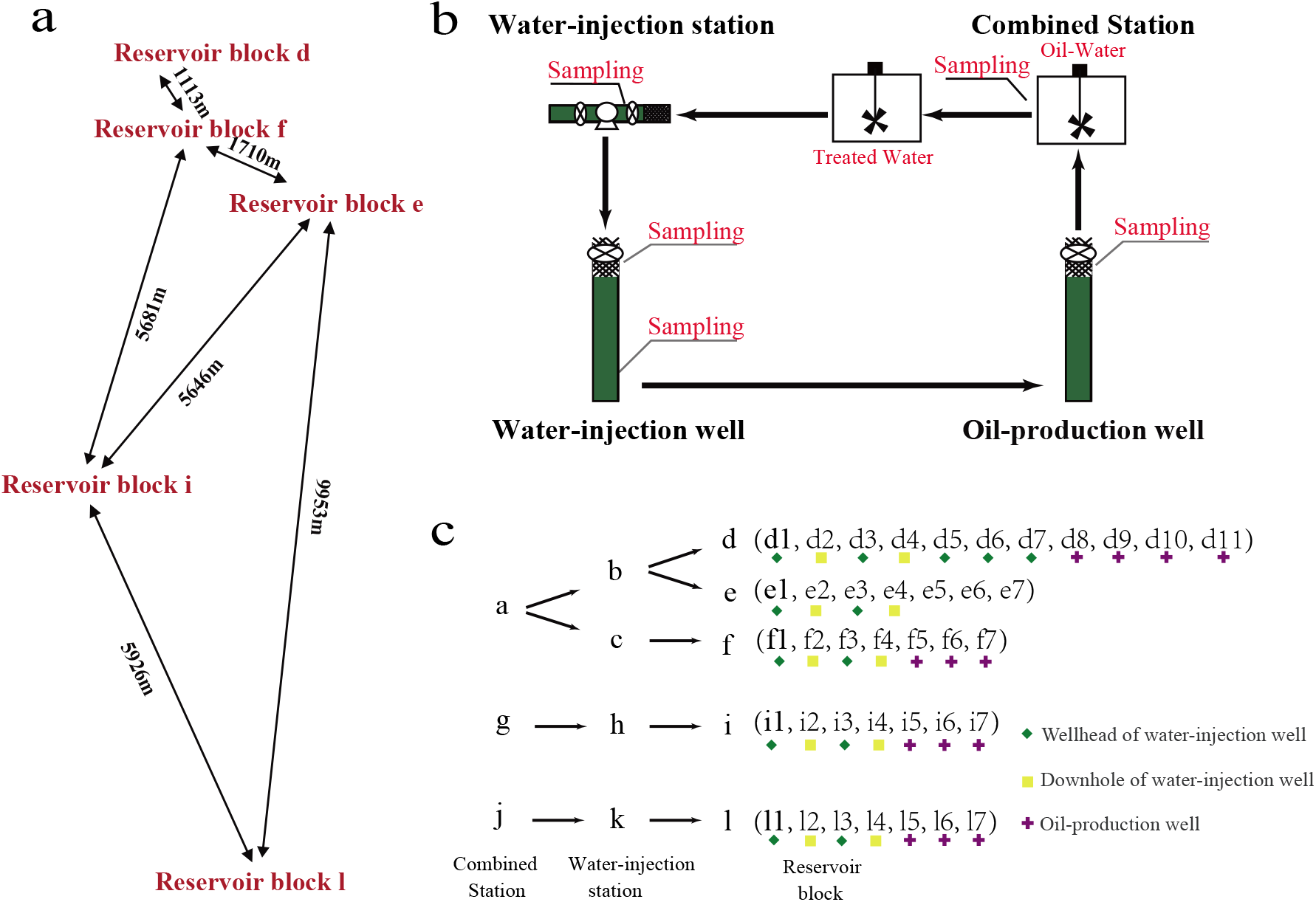
Diagrammatic sketch of the location of the sampled petroleum reservoir block (a), the water-injection facilities and oil-production wells (b and c). The injected water flows from combined injection station (a, g, and j) to water injection station (b, c, h, and k), and then to water-injection wells and oil production wells of reservoir block (d, e, f, i, and l)

In the present work, we investigated the compositions and metabolic profiles of the bacterial and archaeal communities in water injection stations, wellheads, and downhole of water-injection wells and oil-production wells of a water-flooding petroleum reservoir using 16S rRNA gene sequencing, and analyzed the vital influences of environmental variation, microorganisms in injected water, and diffusion-limited microbial transfer in structuring the microbial communities.

## RESULTS

### Microbial community compositions through the oilfield production facilities

After filtering low quality reads and chimeras, a total of 38,102 and 39,729 of bacterial and archaeal sequences on average were obtained for each sample, respectively. The average OTU numbers of the bacterial and archaeal communities were 470 and 67, respectively. The α-diversity indices of the bacterial and archaeal communities in the soil samples were the highest than those in the petroleum reservoir and the water-injection facilities (Fig. 2a and Fig. S1). While similar α-diversity values of the bacterial and archaeal communities was detected between the water-injection stations and the wellheads of the water-injection wells, the samples in the downhole of the water-injection wells showed higher Sobs (546 vs 791, *p* < 0.01) and Shannon (3.89 vs. 4.09, *p* < 0.01) indices for the bacterial communities, and lower values for the archaeal communities (66 vs 72, *p* < 0.01; 1.92 vs. 2.10). A sharp decrease in the Sobs and Shannon indices were observed for both the bacterial (245, *p* < 0.001; 1.95, *p* < 0.001) and archaeal communities (53, *p* < 0.05; 1.61, *p* < 0.001) in the oil-production wells. The Simpson indices were higher in the oil-production wells than in the wellheads and downhole of water-injection wells for both the bacterial (0.33 vs 0.05, *p* < 0.01; 0.33 vs. 0.11, *p* < 0.05) and archaeal communities (0.32 vs. 0.21, *p* < 0.001; 0.32 vs 0.25). Detailed α-diversity indices for the bacterial and archaeal communities are provided in Tables S2 and S3.

**Fig. 2.**
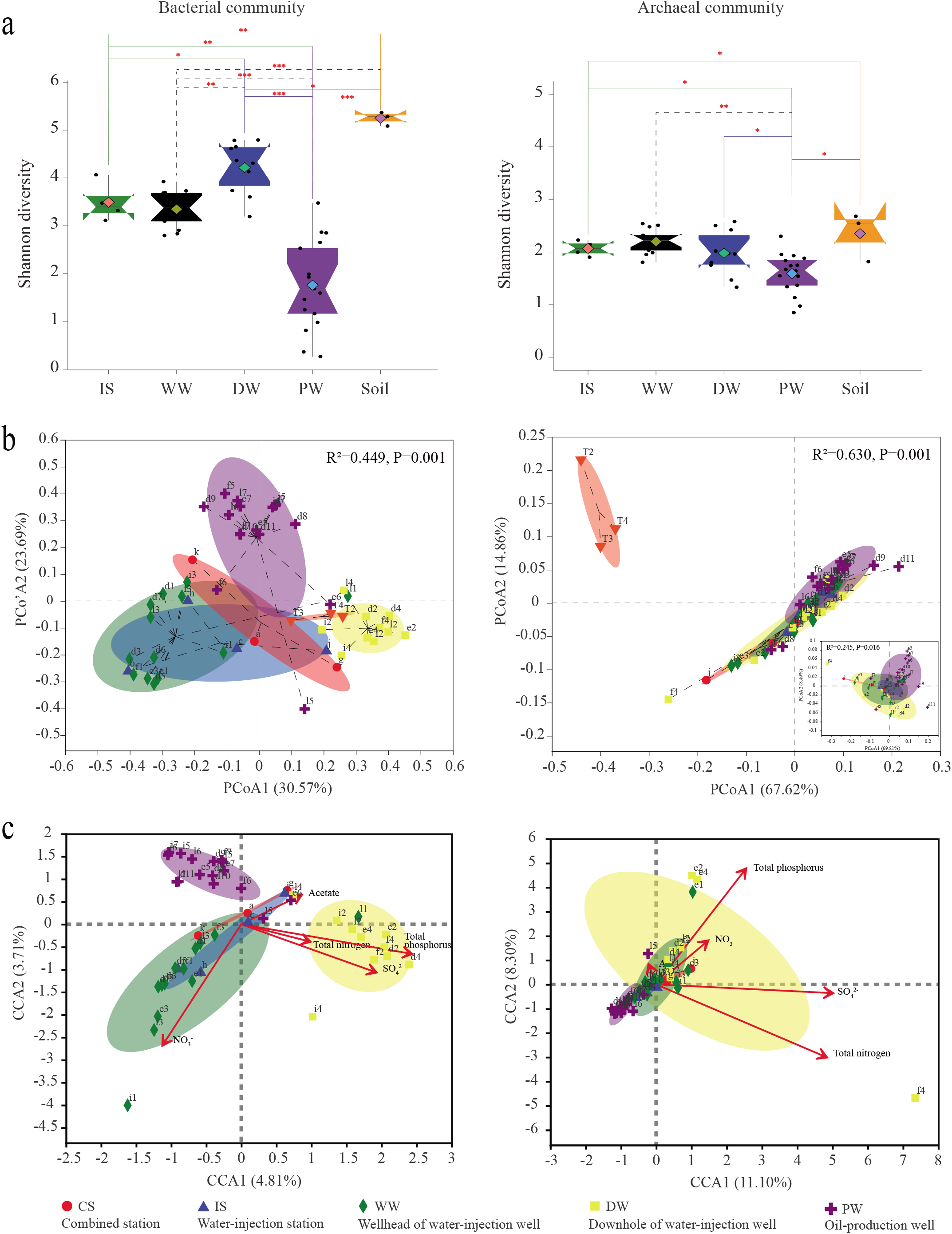
The alpha diversity (a) and beta diversity (b) of the bacterial and archaeal communities from the water-injection facilities and oil-production wells, and (c) the relationships between the community compositions with the environmental variables. PCoA was performed based on Weighted-UniFrac dissimilarity matrixes. Canonical correspondence analysis (CCA) and Monte Carlo permutation test were performed to reveal the correlations between environmental variables and community compositions.

The bacterial communities in the water-injection stations, wellheads, and downhole of the water-injection and oil-production wells showed a similar community composition as shown in the heatmap (Fig. 3a) and cumulative histogram (Fig. S2a). The distributions of the bacterial communities were further visualized via PCoA based on weighted-Unifrac distance matrices (Fig. 2b). The ordination graph suggested that the samples from the water-injection stations and wellheads of the water-injection wells were clustered together, indicating that these locations shared similar community compositions, as confirmed by permutational multivariate analysis of variance (ADONIS; *r*^2^ = 0.083, *p* = 0.221) and similarity analysis (ANOSIM; *r* = 0.115, *p* = 0.223). The microbial communities were dominated by OTUs representing Betaproteobacteria (Comamonadaceae, Alcaligenaceae, and Rhodocyclaceae) and Alphaproteobacteria (Rhodobacteraceae, Sphingomonadaceae, and Burkholderiaceae), followed by Flavobacteria *(Flavobacterium)* and Gamaproteobacteria *(Pseudomonas)* (Fig. 3b and Fig. S3). The samples collected from the wellheads and downhole of the water-injection wells formed distinct clusters in the PCoA plot (Fig. 2b), and the microbial communities in these locations were statistically significant (ADONIS, *r*^2^ = 0.472, *p* = 0.001; ANOSIM, *r* = 0.855, *p* = 0.001). In contrast to the bacterial communities from the wellheads of the water-injection wells, the relative abundances of Clostridia (Clostridiaceae), Deltaproteobacteria (Syntrophaceae, Syntrophorhabdaceae, Desulfobulbaceae, and Desulfovibrionaceae), Anaerolineae (Anaerolineaceae), and Synergistia were significantly higher in the bacterial communities from the downhole of the water-injection wells, where Betaproteobacteria, Alphaproteobacteria, and Flavobacteria were found to diminish markedly (Fig. 3b and Fig. S4). The composition of the bacterial community changed significantly again in the oil-production wells (ADONIS, *r*^2^ = 0.333, *p* = 0.001; ANOSIM, *r* = 0.695, *p* = 0.001). Gammaproteobacteria, Betaproteobacteria, and Epsilonproteobacteria became the most abundant lineages, and *Pseudomonas, Acinetobacter*, *Thauera*, and *Arcobacter* were the dominant genera (Fig. 3b and Fig. S5).

**Fig. 3.**
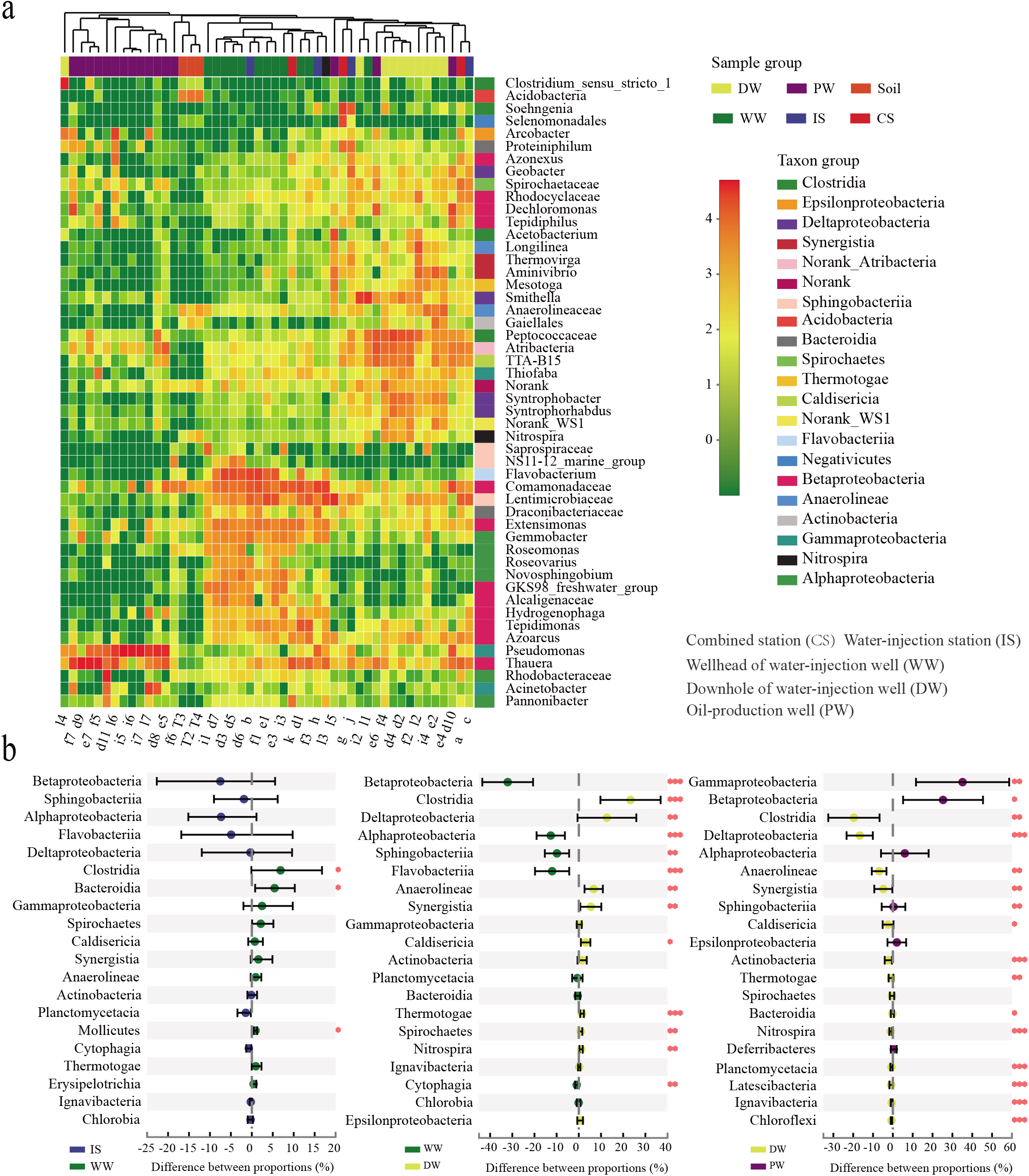
Heatmaps (a) and Wilcoxon rank-sum test (b) showing the distinct distribution of dominant bacterial populations through the water-injection facilities and oil-production wells

Most of the archaeal populations detected existed persistently throughout the water injection and oil production facilities. *Methanosaeta*, *Methanobacterium*, *Methanothermobacter*, *Methanolinea*, *Methanomethylovorans*, *Methanocalculus*, and *Methanoculleus* accounted for the majority of the archaeal sequences in each sample (Fig. 4a and Fig. S2b). However, despite this, *Methanosaeta, Methanobacterium*, and *Methanolinea* were the dominant genera and species in the water-injection facilities, while the oil-production wells were predominated by *Methanosaeta*, *Methanomethylovorans*, and *Methanocalculus*. In addition, significant changes in the archaeal communities were observed between the samples from the downhole of the water-injection and oil-production wells (ADONIS, *r*^2^ = 0.186, *p* = 0.008; ANOSIM, *r* = 0.242, *p* = 0.005). The dominant species *Methanobacterium* and *Methanolinea* were higher in the downhole of the water-injection wells than in the oil-production wells, in which *Methanomethylovorans* (*p* < 0.01), *Methanocalculus* (*p* < 0.01), and *Methanosaeta* were more abundant (Fig. 4b).

**Fig. 4.**
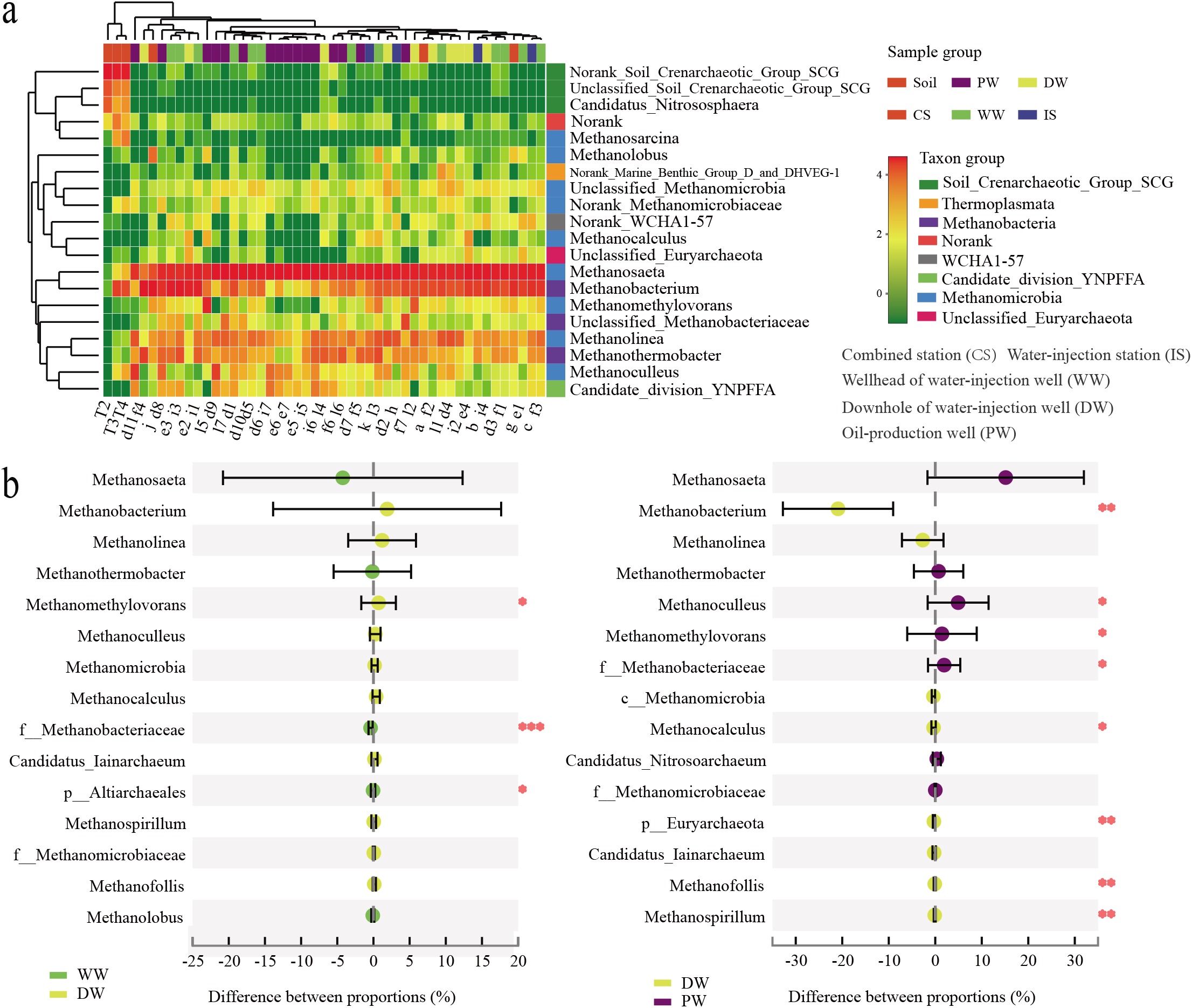
Heatmaps (a) and Wilcoxon rank-sum test (b) showing the distinct distribution of dominant archaeal populations through the water-injection facilities and oil-production wells

### Distinct metabolic profiles of the microbial communities

The metabolic profiles of the bacterial and archaeal communities in the wellheads and downhole of the water-injection and oil-production wells were inferred from 16S rRNA data using Tax4Fun. For the bacterial communities, the majority of the predicted protein sequences annotated with KEGG pathways were clustered into metabolism (56.42-63.66%), environmental information processing (15.62-22.42%), genetic information processing (9.08-14.25%), and cellular processes (3.97-7.42%). Significant differences were observed in the aforementioned pathways among the wellheads and downhole of the water-injection, and oil-production wells as illustrated in Fig. 5 and Fig. S6-8a. The sequences related to the biosynthesis of other secondary metabolites were found to have the highest abundance in the wellheads of the water-injection wells (Fig. 5a)(*p* < 0.05). In addition, the relative abundance of the sequences related to amino acid metabolism, xenobiotic biodegradation and metabolism, and lipid metabolism were higher than those of the downhole of the water-injection wells (Fig. 5a) (*p* < 0.05), while amino acid metabolism, carbohydrate metabolism, translation, nucleotide metabolism, replication and repair, and glycan biosynthesis and metabolism had higher abundances than those of the oil-production wells (Fig. 5a) (*p* < 0.05). In the downhole of the water-injection wells, the relative abundance of sequences associated with energy metabolism, translation, nucleotide metabolism, and glycan biosynthesis and metabolism were higher than those of the wellheads of the water-injection and oil-production wells (Fig. 5a) (*p* < 0.05). Compared with the wellheads and downhole of the water-injection wells, the sequences associated with cell motility (bacterial chemotaxis), cellular community (biofilm formation), and signal transduction (two-component system) were more abundant in oil-production wells (Fig. 5a) (*p* < 0.05). In addition, the sequences related to xenobiotic biodegradation and metabolism, lipid metabolism, metabolism of terpenoids and polyketides, amino acid metabolism, and membrane transport were higher than those of the downhole water-injection wells (Fig. 5a) (*p* < 0.05).

**Fig. 5.**
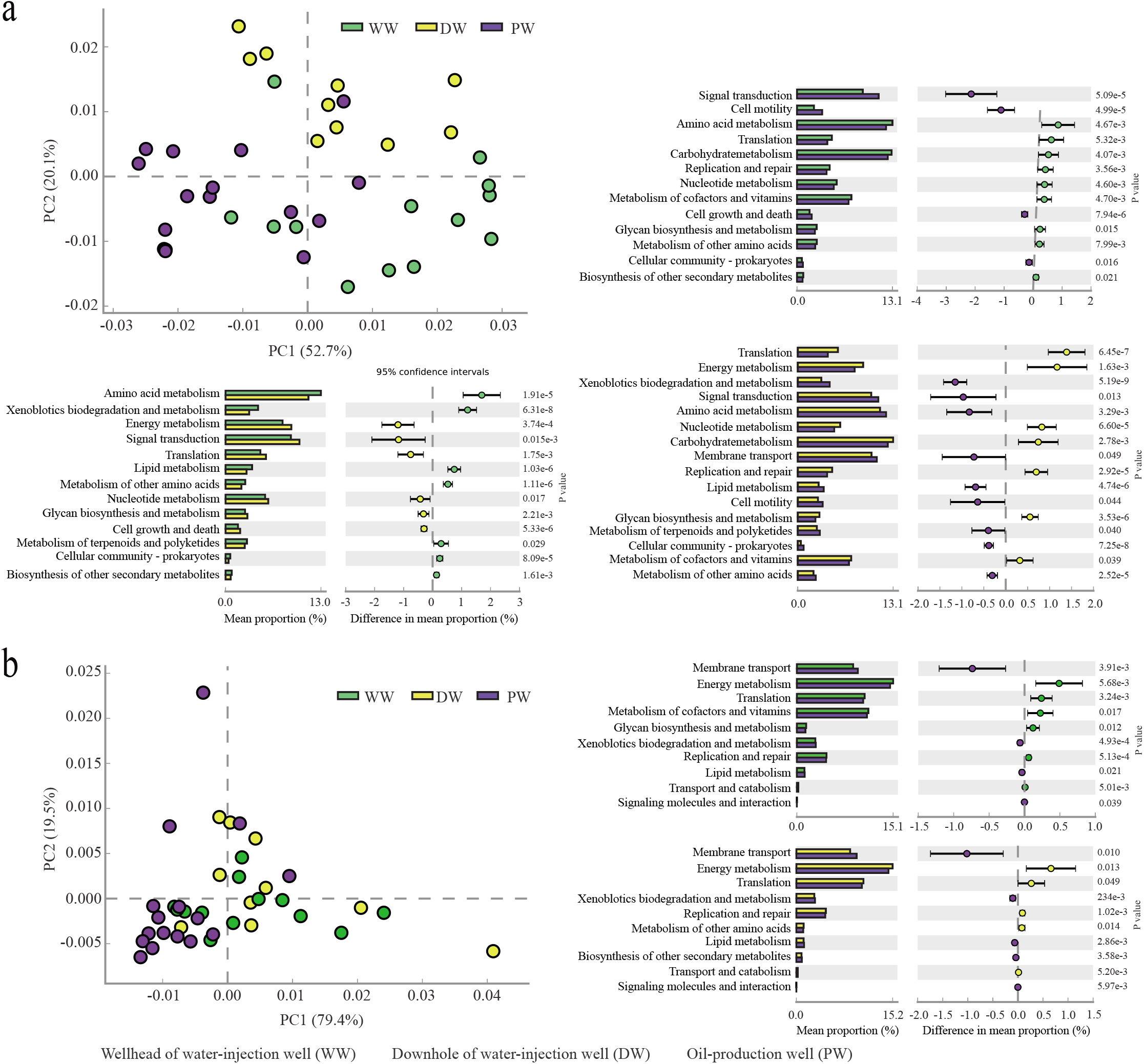
Distinct metabolic profiles and the statistically significant differences among the bacterial (a) and archaeal (b) communities through wellheads and downhole of the water-injection wells and the oil-production wells. The ordination graph showed that the samples with similar metabolic profiles were clustered together, otherwise, formed distinct clusters. The bar charts show the differences between the proportions of sequences in each group with a confidence interval of 95%

For the archaeal communities, the sequences clustered into metabolism, environmental information processing, genetic information processing, and cellular processes accounted for 62.97-63.43%, 8.05-13.13%, 19.18-20.51%, and 1.38-2.70%, respectively. Significant differences were observed in the pathways between the water-injection and oil-production wells (Fig. 5b and Figs. S6-8b). In the wellheads and downhole of water-injection wells, the relative abundance of the sequences related to energy metabolism, translation, metabolism of cofactors and vitamins, glycan biosynthesis and metabolism, and replication and repair were higher than those of the oil-production wells (Fig. 5b) (*p* < 0.05). However, the sequences associated with membrane transport (bacterial secretion system), lipid mechanism, and xenobiotic biodegradation and metabolism were found to have higher abundances in the oil-production wells (Fig. 5b) (*p* < 0.05).

### Distinct network patterns of the bacterial communities

Co-occurrence network analysis was used to assess the interactions of microbial populations within the microbial communities inhabiting the water-injection and oil-production wells. The complexities of the networks were compared based on the number of nodes, edges, average degrees, clustering coefficient, scale-free, and modularity (Fig. 6). The phylogenetic molecular ecological networks were constructed with similarity thresholds of 0.81, 0.98, and 0.81 for the wellheads and downhole of the water-injection and oil-production wells, respectively. However, a greater number of nodes and links were observed in the downhole network of the water-injection wells, followed by the wellheads of the water-injection wells, followed by the oil-production wells. Compared with the wellhead network (R^2^ = 0.463) and the network of the oil-production wells (R^2^ = 0.197), the downhole network was closely fitted with the power-law model (R^2^ = 0.863), representing a scale-free network, in which few nodes in the network have a large number of neighbors and most nodes have few neighbors. There was a higher average degree and average clustering coefficient and a lower centralization of degree and betweenness of the nodes in the wellhead network. The modularity and number of modules were higher in the downhole and oil-production well networks than in the wellhead network. In addition, there were more positive correlations in the networks of the wellheads and downhole of the water-injection wells, while more negative correlations were observed in the oil-production wells.

**Fig. 6.**
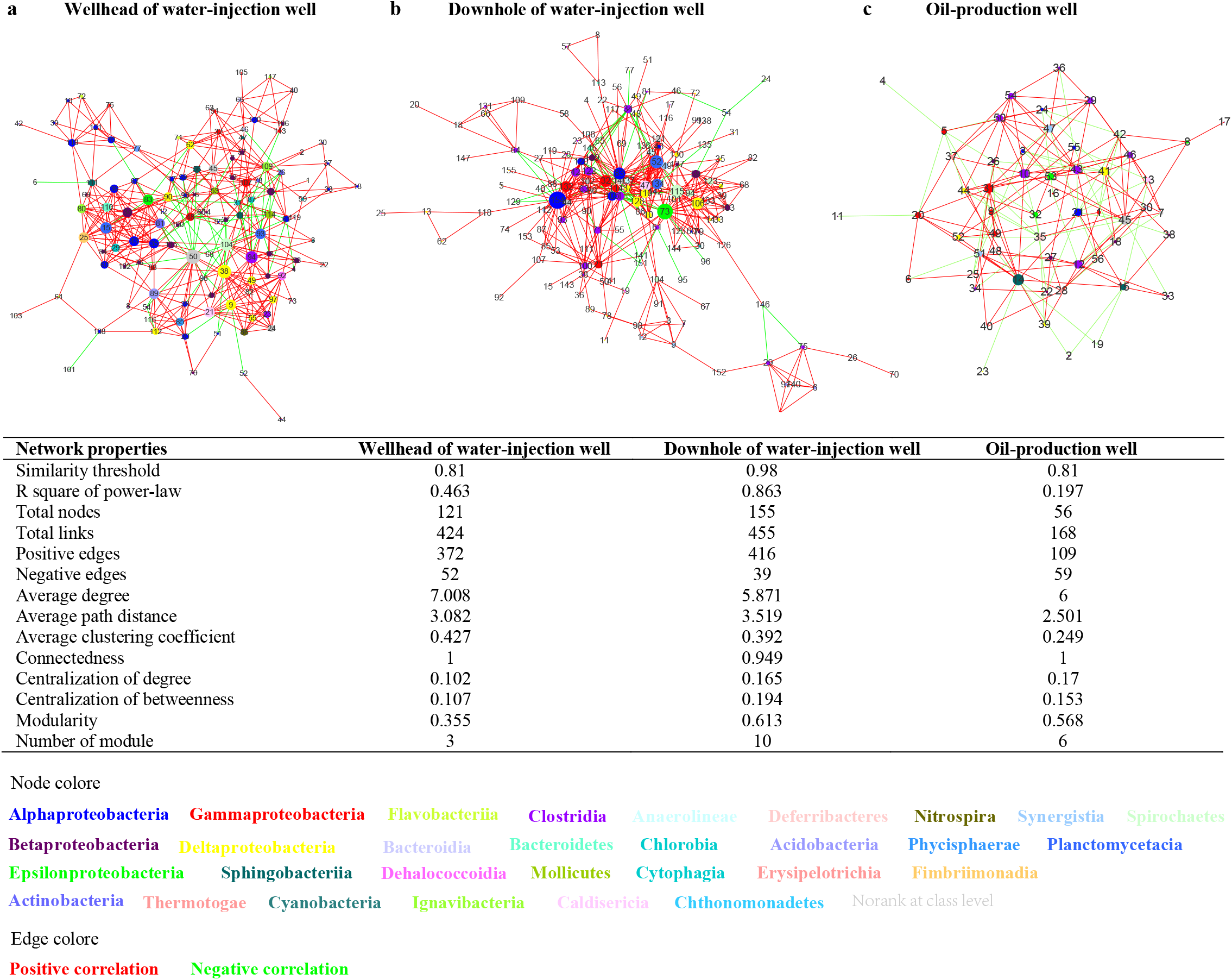
Co-occurrence networks and topological properties of the bacterial and archaeal communities through wellheads (a) and downhole (b) of the water-injection wells and the oil-production wells (c). Nodes are colored according to microbial class, and the nodes with a larger size show the potential keystone OTUs. The taxonomic information for the numbered nodes and the potential keystone OTUs is listed in Table S4. Edges indicate correlations among nodes, and the red and green edges represent positive and negative correlations, respectively

The potential keystone taxa in each network were screened (Table S4), including those that act as connectors, module hubs, network hubs, and those with low betweenness and/or high degree (33–36). More nodes were assigned to Betaproteobacteria, Alphaproteobacteria, Flavobacteriia, and Sphingobacteriia in the wellhead network. The bacterial keystone taxa were mainly from Flavobacteriaceae, Commamonadaceae, Lentimicrobiaceae, Syntrophaceae, Pseudomonadaceae, Sphingomonadaceae, Rhodobacteraceae, and Alcaligenaceae, including the dominant genera *Flavobacterium, Novosphingobium, Gemmobacter, Roseovarius, Smithella*, and *Pseudomonas*. There were more nodes from Clostridia, Deltaproteobacteria, Anaerolineae, Nitrospira, Betaproteobacteria, and Synergistia in the downhole network. The keystone taxa mainly belonged to Clostridiaceae, Anaerolineaceae, Syntrophaceae, Rhodocyclaceae, Desulfovibrionaceae, and Desulfobulbaceae, including the dominant genera *Clostridium*, *Nitrospira*, *Syntrophus*, *Smithella*, *Desulfovibrio*, and *Desulfobulbus*. In the network of the oil-production wells, the nodes were mainly from Gammaproteobacteria, Betaproteobacteria, Epsilonproteobacteria, and Deltaproteobacteria, and the OTUs that were assigned to Rhodocyclaceae and Syntrophaceae were the main keystone taxa, including dominant genera *Thauera*, *Azoarcus*, *Dechloromonas*, and *Smithella*. This is also reflected in the differences observed in the overall community compositions among oilfield production facilities.

### Environmental selection on the microbial communities

To observe changes in the environmental variables of water-injection facilities and oil-production wells, NMDS and ADONIS analyses were performed using the contents of acetate, NO_3_^-^, SO_4_^2-^, total nitrogen, and total phosphorus. As shown in the NMDS plot, the samples collected from similar locations were clustered together (Fig. S9). No significant differences in the environmental factors were observed between the water-injection stations and the wellheads of the water-injection wells. However, significant differences were observed among the wellheads and downhole of the water-injection and oil-production wells (Table S5). The wellheads of the water-injection wells had a higher NO_3_^-^ concentration (8.33 mg/L), which was found to decrease significantly in the downhole of the water-injection wells (2.66 mg/L, ADONIS, R^2^ = 0.41, p ⩽ 0.01) and the oil-production wells (0.90 mg/L, ADONIS, R^2^ = 0.41, p ⩽ 0.01). The concentration of SO_4_^2-^ (32.10 mg/L) in the downhole of the water-injection wells was over 2.5-fold that in the wellheads of the water-injection wells (12.01 mg/L) and 6-fold that in the oil-production wells (5.18 mg/L). The concentration of phosphorus (29.64 mg/L) in the downhole of the water-injection wells was over 10-fold that in the wellheads of the water-injection wells (2.83 mg/L) and the oil-production wells (2.88 mg/L). There were no significant changes in the acetate concentrations, which averaged 11.24-11.69–mg/L in the water-injection and oil-production wells (Table S5).

The effects of environmental variations on the spatial distributions of the bacterial and archeal communities were further analyzed via the Mantel test, ADONIS, and CCA analysis. The Mantel test showed significant correlations between the environmental variables and microbial community compositions through the water-injection facilities and oil-production wells (Table S6). Significant correlations were observed between the bacterial communities and the contents of acetate and NO_3_^-^ (Table S6; Mantel, p = 0.011 and 0.007, respectively), and between the archaeal communities and NO_3_^-^ content (Table S6; Mantel, p = 0.008) in the water-injection pipelines consisting of combined stations, water-injection stations, and the wellheads of water-injection wells. For the wellheads and downhole of the water-injection and oil-production wells, significant correlations were observed between the bacterial communities and the total phosphorus, NO_3_^-^, and SO_4_^2-^ contents (Table S6; Mantel, p = 0.001), and between the archaeal communities and the total nitrogen content (Table S6; Mantel, p = 0.045). ADONIS showed that total phosphorus and SO_4_^2-^ can effectively explain the changes in the compositions of the bacterial community in the wellheads and downhole of the water-injection wells, and in the downhole of the water-injection and oil-production wells (Table S7). For the archaeal communities, significant correlations between community composition and total phosphorus and SO_4_^2-^ were observed for the downhole of the water-injection and oil-production wells (Table S6). Despite the high correlation, CCA analysis indicated that the environmental variables could only explain 14.8% of the bacterial community changes, and 24.8% of the archaeal community changed through the injection-production facilities (Fig. 2c and Table S8).

### Diffusion-limited microbial transfer in the water-injection pipelines and oil-bearing strata

The persistent OTUs were analyzed to elucidate a potential transfer of microorganisms in the water-injection pipelines and oil-production wells. As shown in Venn diagrams, a large number of persistent OTUs were detected through the water-injection facilities and oil-production wells (Fig. S10). Correlation analysis indicated that the water-injection stations and the wellheads of the water-injection wells harbored a large number of shared OTUs with similar relative abundances (Fig. 7a; Pearson correlation coefficient (*r*) = 0.78, p < 0.001 for bacteria and *r* = 0.97, p < 0.001 for archaea). A large number of shared OTUs with different relative abundances were detected in the wellheads and downhole of water-injection wells (Fig. 7a, *r* = 0.78, p < 0.001 for bacteria and *r* = 0.97, p < 0.001 for archaea). There were also substantial OTUs with different relative abundances in the downhole of the water-injection and oil-production wells (Fig. 7a, *r* = 0.78, p < 0.001 for bacteria and *r* = 0.97, p < 0.001 for archaea). As shown in Fig. 7b, 51.1-74.9% of the OTUs detected in the downhole of the water-injection wells were detected in the wellheads of the water-injection wells, and 8.2-43.9% of the OTUs from the oil-production wells appeared in the downhole of the water-injection wells. In addition, the proportions of the shared OTUs in the oil-production wells showed strong correlations with the formation permeability of the oil-production wells. In fact, in blocks d, f, and i, the correlation coefficient reached 0.961 (Fig. 7b; p < 0.01). The correlation of the shared OTUs with the formation permeability of the oil-production wells was not clear in blocks e and l (Fig. 7b).

**Fig. 7.**
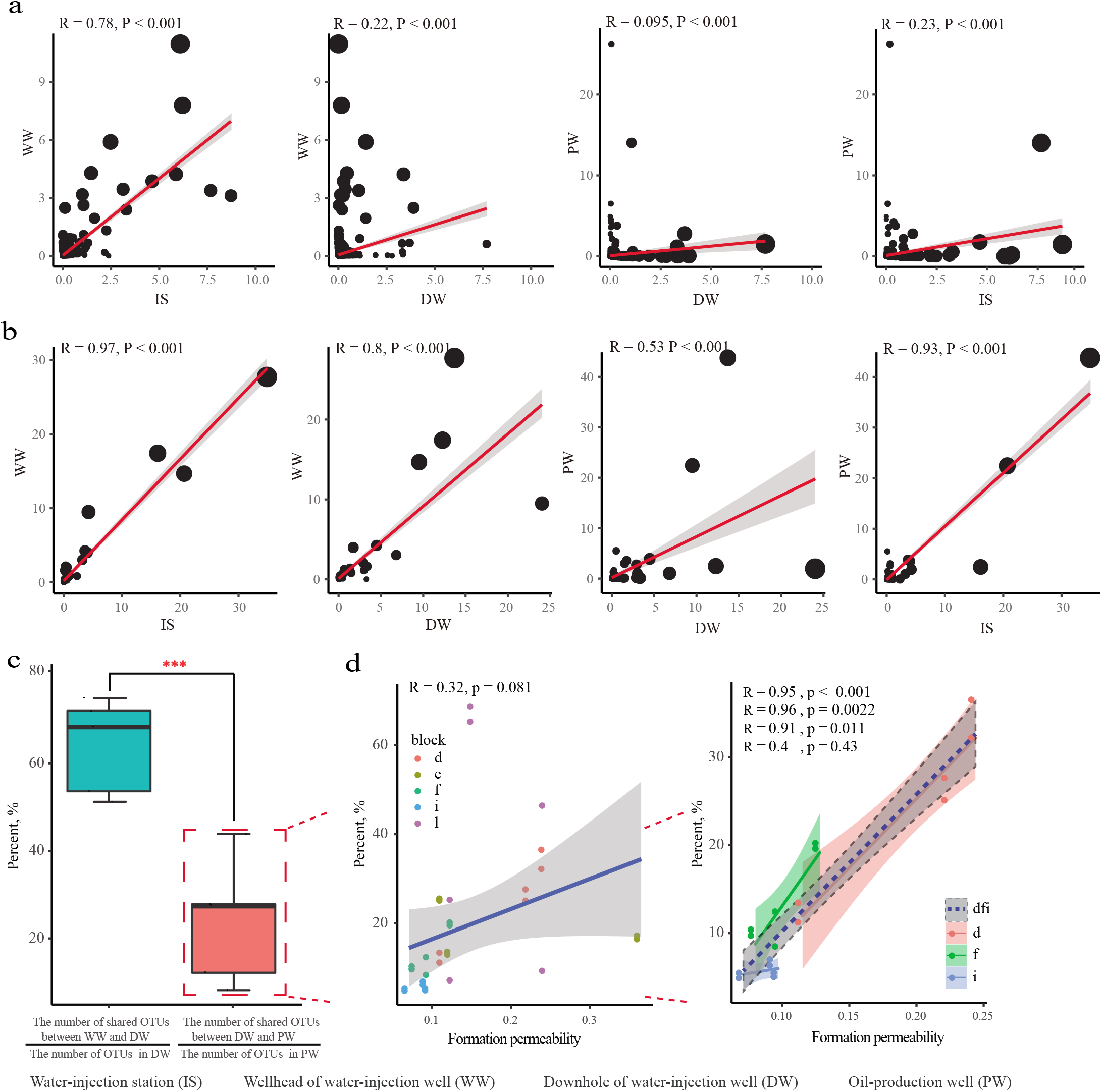
Correlations of the bacterial (a) and archaeal (b) communities inhabiting in the water-injection facilities and oil-production wells, (c) the distribution of shared bacterial OTUs between wellheads and downhole of the injection wells, and downhole of the injection wells and oil-production wells, and (d) the correlations with stratal permeability of the oil production wells.

Moreover, diffusion-limited microbial transfer in oil-bearing porous medium was tested in oil-bearing cores using strain SG-rfp marked by a red fluorescent protein-encoding gene (37). For the cores with a permeability of 1.405 μm^2^, a bright fluorescent signal was observed in the effluent when 1 pore volume (PV) of the displacing fluid containing RFP-labeled *Pseudomonas aeruginosa* was injected. The maximum fluorescence was observed when 2 PV displacing fluid was injected (Fig. S11). For the cores with a permeability of 0.203 μm^2^ , a fluorescent signal was clearly observed in the effluent when 2 PV of displacing fluid was injected. The maximum fluorescence was observed until 15 PV displacing fluid was injected (Fig. S11). These results indicate that microorganisms in injected water can migrate through oil-bearing porous medium with the water flow, and that the formation permeability imposed significant limitations on microbial migration.

## DISCUSSION

Microbial-enhanced oil recovery in production is well known for the involvement of microorganisms and their metabolites under a wide spectrum of oil reservoir types (5, 8, 9). To substantiate the process of microbiologically improving oil production is more effective, a clear understanding of the distribution of microbial communities in oilfield production facilities is necessary. As a result, this study systematically investigated the composition and metabolic characteristics of microbial communities in water-injection pipelines and oil-production wells of a water-flooding petroleum reservoir, and revealed the roles of environmental variation, microorganisms in injected water, and diffusion-limited microbial transfer in determining the structure of the microbial communities.

The compositions of the microbial community in oilfield production facilities show spatial specificity. Microbial community α-diversity reflects the number of species in a local homogeneous habitat. In the oilfield production facilities, we found that the α-diversity of both the bacterial and archaeal communities was significantly lower than that found in the ground surface soil. The downhole of the water-injection wells had a higher community α-diversity, while a lower community α-diversity was observed in the oil-production wells. In petroleum reservoirs, both the bacterial and archeal community α-diversity is influenced by extreme environmental conditions and oil-production processes, such as high temperature, hypersalinity, and oil recovery methods (25). This phenomenon may be explained by the adaptability and metabolic types of the microorganisms (see analysis of metabolic profiles). The water-injection stations and the wellheads of the water-injection wells were found to contain similar bacterial communities, dominated by Betaproteobacteria (Comamonadaceae, Alcaligenaceae, and Rhodocyclaceae), Alphaproteobacteria (Rhodobacteraceae, Sphingomonadaceae, and Burkholderiaceae), Flavobacteria (Flavobacteriaceae), including dominant genera *Tepidimonas*, *Extensimonas*, *Hydrogenophaga*, *Flavobacterium*, *Thauera*, *Smithella*, *Novosphingobium*, *Gemmobacter*, *Roseovarius*, *Azoarcus*, *Azovibrio*, and *Rhodobacter*. Most of these populations were comprised of aerobic or facultative anaerobic. The species of the genus *Tepidimonas* (Comamonadaceae) are generally strictly aerobic and chemolithoheterotrophic (38). *Hydrogenophaga* (Comamonadaceae) species are hydrogen-oxidizing bacteria that are able to ferment organic acids (39). *Thauera* (Rhodocyclaceae) has been described as isopropanol, acetone, and aromatic hydrocarbons, and is the main contributor to the mitigation of biological souring in oil reservoirs (12, 40). Some species from *Smithella* have been described as anaerobic propionate-degrading syntrophs (41). Some species of *Azoarcus* (Rhodocyclaceae) are able to fix nitrogen (42). *Rhodobacter* species (Rhodobacteraceae) show a wide range of metabolic capabilities, including photosynthesis, lithotrophy, aerobic and anaerobic respiration, nitrogen fixation, and the synthesis of tetrapyrroles, chlorophylls, heme, and vitamin B12 (43). The downhole of the water injection wells was dominated by Clostridia (Clostridiaceae), Deltaproteobacteria (Syntrophaceae, Syntrophorhabdaceae, Desulfobulbaceae, and Desulfovibrionaceae), Anaerolineae (Anaerolineaceae), Synergistia, and Nitrospira, including dominant genera *Clostridium*, *Syntrophus*, *Smithella*, *Desulfovibrio*, *Desulfobulbus*, *Longilinea*, *Aminiphilus*, *Thermovirga*, and *Nitrospira*. *Clostridium* is a genus of a group of strictly anaerobic Gram-positive bacteria, and has been widely used for the production of organic acids, organic solvents, and enzymes, such as acetone, butanol, 1,3-propanediol, ethanol, butanol, acetic acid, and biohydrogen (44). *Syntrophus* species were reported to be able to degrade benzoate into acetate and H_2_, which were subsequently converted to methane by *Methanosarcina* and *Methanoculleus*, and the direct interspecies electron transfer of *Desulfovibrio* and *Methanosarcina* (45). *Desulfovibrio* and *Desulfobulbus* are commonly detected sulfate-reducers that reduce sulfates to hydrogen sulfide in petroleum reservoirs (46). Anaerobic amino-acid-degrading *Thermovirga* has been previously isolated from a North Sea oil well (47). The oil-production wells were dominated by Gammaproteobacteria (Pseudomonadaceae), Betaproteobacteria (Rhodocyclaceae, Hydrogenophilaceae), Epsilonproteobacteria (Campylobacteraceae), and Deltaproteobacteria (Geobacteraceae), including facultative anaerobic dominant genera *Pseudomonas*, *Thauera*, *Hydrogenophaga*, *Acinetobacter*, *Atribacteria*, *Arcobacter*, *Acetobacterium*, and *Geobacter*. Many strains of *Pseudomonas* and *Acinetobacter* are capable of utilizing hydrocarbons and producing biosurfactants (6, 48, 49). *Geobacter* plays an important role in electron exchange by direct interspecies electron transfer (1). Most of the detected archaeal populations persistently existed throughout the oilfield production facilities. It is worth noting that *Methanosaeta*, *Methanobacterium*, and *Methanolinea* predominated in the water-injection facilities, while the oil-production wells were dominated by *Methanosaeta*, *Methanomethylovorans*, and *Methanocalculus*. *Methanosaeta* is an acetoclastic methanogen that uses only acetate in methane production, while *Methanobacterium*, *Methanolinea*, *and Methanocalculus* are methylotrophic and hydrogenotrophic methanogens (50). *Methanomethylovorans* is a methylotrophic methanogen that is able to grow on dimethyl sulfide and methanethiol (51). It is not difficult to see that microorganisms inhabiting petroleum reservoirs maintain a close association with each other and show a high metabolic potential for hydrocarbon degradation, sulfate reduction, nitrate/nitrite reduction, and methanogenesis.

Associating community compositions with functional predictions enabled us to decipher the potential ecological traits of the microbial communities found throughout oilfield production facilities. Significant differences in metabolism, environmental information processing, genetic information processing, and cellular processes pathway were observed among the wellheads and downhole of the water-injection and oil-production wells. Pathways associated with the biosynthesis of other secondary metabolites, amino acid metabolism, xenobiotic biodegradation and metabolism, and lipid metabolism likely played more important roles in the wellheads of the water-injection wells. By contrast, the downhole of the water-injection wells showed more activity in pathways associated with energy metabolism, translation, nucleotide metabolism, and glycan biosynthesis and metabolism. These findings suggest that the growth and metabolism of microorganisms in the downhole of the water-injection wells were more active. In the oil-production wells, cell motility (bacterial chemotaxis), cellular community (biofilm formation), and signal transduction (two-component system) were found to be more active. In addition, xenobiotic biodegradation and metabolism, metabolism of terpenoids and polyketides, amino acid metabolism, and membrane transport also played important roles. Bacteria usually use chemotaxis to position themselves within the optimal portion of their habitats to approach specific chemical attractants and avoid repellent ligands (52). Both bacteria and archaea are capable of forming biofilms, which often benefit the survival of microorganisms in the presence of environmental stresses, such as low or high pH and toxic chemicals, and facilitate horizontal gene transfer and syntrophy with other microorganisms (53). Due to the extreme environment in oil reservoirs, microbial populations most likely tend to metabolize substrates in the form of synergetic metabolism and mutualism. The abundance of two-component systems suggests that the microorganisms in the oil-production wells are likely to have suffered more extreme environmental stresses, particularly nutritional deficiency and oxygen limitation. The two-component systems also regulate a variety of physiological behaviors of microorganisms, such as motility, chemotaxis (54), spore formation (55), and biofilm formation (56).

The microbial community composition throughout oil-production facilities was closely correlated with to the nutrients available. Mantel test, ADONIS, and CCA analysis revealed significant correlations between the total phosphorus, NO_3_^-^, and SO_4_^2-^ contents and the compositions of the microbial communities throughout the oilfield production facilities. It is worth noting that the phosphorus and SO_4_^2-^ contents in the downhole of the water-injection wells were far greater than those in the wellheads of the water-injection and oil-production wells. This is consistent with higher activity in the growth and metabolism of microorganisms in the downhole of the water-injection wells. With the increase in the SO_4_^2-^ content, the abundance of sulfate-reducers, such as *Desulfovibrio* and *Desulforhabdus*, also significantly increased. The accumulation of sulfate and other nutrients (especially phosphate) in the downhole of the water-injection wells may be interpreted as chemical deposition under the action of formation brines and the interception role because of the sieve effect of oil-bearing strata. Despite demonstrating a high correlation, the nutrient distribution only partially explained the changes in the bacterial and archeal communities throughout the injection-production facilities. Oxygen levels, temperature (13, 22, 57), salinity (23, 24, 58), and pH (25, 59) are other major determinants of microbial community composition. As the results of the ANOSIM and ADONIS analyses reveal, strong relationships were observed between sampling sites and microbial community compositions. Combined with the distribution of dominant microbial populations throughout the injection-production facilities, oxygen levels are likely to play a crucial role in determining community composition.

The water-flooding process seems to continually inoculate reservoirs with exogenous microorganisms. However, whether the microorganisms in injected water can pass through oil-bearing strata and reach oil-production wells, and the influence of these microorganisms on subsurface microbial communities, remains unclear. The low permeability of oil-bearing strata inevitably exerts a significant influence on microbial diffusion in oil reservoirs. Lenchi et al. reported that the bacteria associated with water injected into oil reservoirs were not retrieved from oil-production waters (60). Further research has found that a large number of shared OTUs were detected in water-injection wells and adjacent oil-production wells, with aerobic populations often appearing in oil-production wells (17, 18, 31, 32). It seems that microorganisms on the ground may migrate or be brought into oil reservoirs during the oil production process. Ren et al. recently suggested that the transportation of injected bacteria in oil-bearing strata was impacted by the varied permeability from water-injection wells to adjacent oil-production wells (30). In the present study, a large number of OTUs with similar relative abundances were observed simultaneously in the water-injection stations and the wellheads of the water-injection wells, and greater rations of shared OTUs were observed between the wellheads and downhole of the water-injection wells than those between the downhole of the water-injection wells and the oil-production wells. These findings imply that microorganisms may migrate in water flowing in water-injection pipelines and oil-bearing strata. Furthermore, the rations of shared OTUs detected in the downhole of the water-injection and oil-production wells showed strong correlations with the corresponding formation permeability, highlighting the influence of geographic isolation on microbial transfer in oil-bearing strata. This phenomenon was further demonstrated using the core-flooding test. Although microorganisms can be brought into oil-bearing strata even in oil-production wells, their influence on the subsurface microbial communities is closely related to their adaption and growth in new environments.

In this study, we revealed the spatial distribution of microbial communities in the oil-production facilities of a water-flooding petroleum reservoir. Our results indicate that environmental variation and microorganisms in injected water are the determinants of the structure of microbial communities in water-injection facilities, while the determinants in oil-bearing strata are environmental variation and diffusion-limited microbial transfer. These findings provide further insights into the distribution of microbial communities in oil-production facilities and could have profound consequences on the use of reservoir microorganisms for improving the oil production process. However, future studies will need to quantify the effects of these factors on structuring the microbial communities in oil-production facilities.

## MATERIALS AND METHODS

### Sampling sites and samples collection

A water-flooding petroleum reservoir of the Daqing Oilfield in the northeast China was chosen for this study. The temperature of the reservoir was approximately 45°C. Samples were collected from combined stations a, g, and j, corresponding to oil-water separation and water treatment before water re-injection into the well, water-injection stations b, c, h, and k, for transferring water to wellheads and downhole of water injection wells, and oil-production wells of blocks d, e, f, i, and l (Fig. 1). The water of water-injection stations b and c was from combined station a. The injected water flowed from b into the water-injection wells of blocks d and e, and flowed from c into the water-injection wells of block f. The injected water at injection station h was from combined station g and flowed into the water-injection wells of block i. The injected water of injection station k was from combined station j and flowed into the water-injection wells of block l. The average permeability of block d is 0.3 μm^2^, ranging from 0.197 μm^2^ to 0.5 μm^2^. The average permeability of block f is 0.289 μm^2^, ranging from 0.125 μm^2^ to 0.811 μm^2^. The average permeability of block e is 0.069 μm^2^, ranging from 0.07 μm^2^ to 0.128 μm^2^. The average permeability of block i is 0.099 μm^2^, ranging from 0.071 μm^2^ to 0.121 μm^2^. The average permeability of the block l is 0.204 μm^2^, ranging from 0.028 μm^2^ to 0.245 μm^2^.

The samples from the downhole of the water-injection wells were obtained by water backflow; that is, injected water flowed upward through the well-hole under pressure. The other samples were taken through the reserved sampling valves of the water-injection and oil-production facilities. The samples collected filled 10-L sterilized plastic buckets, which were tightly sealed with screw caps, and immediately transported to the laboratory for DNA extraction and chemical analysis. The samples were numbered with the sampling sites, such as d1 and d2, which represent the samples collected from the wellheads and downhole of the water-injection well d1 in block d. In addition, three soil samples labeled as T1, T2, and T3 were collected from the ground soil (at a depth of 5 cm) of the water injection wells d1, d3, and d5, respectively.

### Chemical analysis and DNA extraction

The concentrations of acetate, NO_3_^-^, and SO_4_^2-^ in the water samples were determined using an ion chromatograph (DIONEX ICS-1000) equipped with a Shim-pack IC-C3 column. Total nitrogen and phosphorus were analyzed according to “HJ 636-2012 Water quality - Determination of total nitrogen - Alkaline potassium persulfate digestion UV spectrophotometric method” and “GB 11893-1989 Water quality - Determination of total phosphorus - Ammonium molybdate spectrophotometric method”, respectively. Detailed data are listed in Table S1. Microbial cells were collected from to 2-3 L of water samples by centrifugation at 12,000 × *g* and 4°C for 20 min in a high-speed centrifuge (Beckman, USA). Total genomic DNA was extracted using AxyPrep™ Genomic DNA Miniprep Kit (Axygen, USA) combined with bead shaker treatment, as previously described (5).

### 16S rRNA sequencing and bioinformatics analysis

The bacterial and archaeal 16S rRNA genes were amplified using the universal prokaryotic primers 515f (5’-GTG CCA GCM GCC GCG GTA A-3’), 907r (5’-CCG TCA ATT CMT TTR AGT TT-3’), 524f (5’-TGY CAG CCG CCG CGG TAA-3’), and 958r (5’-YCC GGC GTT GAV TCC AAT T-3’), respectively. PCR amplicons were paired-end sequenced (2× 250 bp) on an Illumina MiSeq platform, according to the standard protocol (Majorbio Bio-Pharm Technology Co., Ltd, Shanghai, China). Raw fastq files were demultiplexed and quality-filtered using QIIME2 (61). The sequences were assigned to operational taxonomic units (OTUs) at a 97% sequence similarity level using the UPARSE pipeline (62). The representative sequence sets were aligned and given a taxonomic classification by RDP (63) against the SILVA Small Subunit rRNA database at an 80% confidence threshold. The α-diversity, including observed OTUs, Chao1, Shannon, and Simpson indices, was calculated based on the OTUs. The community β-diversity was estimated based on weighted-UniFrac dissimilarity between samples.

Tax4Fun (64) was used to predict the functional profiles of the microbial communities based on the 16S rRNA data obtained on Majorbio Cloud Platform (www.majorbio.com). Tax4Fun transforms the SILVA-based OTU classification into a taxonomic profile of identical or closely-related genomes in the Kyoto Encyclopedia of Genes and Genomes (KEGG) database. These taxonomic profiles were converted into artificial metagenomes/metatranscriptomes by incorporating the functional data calculated from the genomes of each KEGG organism. The statistical significance of differentially abundant functional categories was tested using Statistical Analysis of Metagenomic Profiles (STAMP) software (65). To explore potential interactions among the microbial populations, co-occurrence network analyses were carried out based on the Pearson correlation between OTUs using Molecular Ecological Network Analyses Pipeline (MENAP) (33). Poorly represented OTUs (i.e. those existing in fewer than 50% of the samples and had less than 0.05% average relative abundance in each group) were removed from the network analyses. To describe the topology of the resulting network, average node connectivity, average path length, diameter, cumulative degree distribution, clustering coefficient, and modularity were calculated. The constructed networks were visualized using Cytoscape version 3.7.2 (66).

### Statistical analysis

Changes in the community α-diversity of the water-injection and oil-production facilities were analyzed using analysis of variance (ANOVA). To visualize the relationships of the microbial communities, principal coordinates analysis (PCoA) was performed based on weighted-UniFrac dissimilarity matrices. To determine the significant differences in the microbial β-diversity of the water-injection and oil-production facilities, permutational multivariate analysis of variance (ADONIS) and similarity analysis (ANOSIM), depending on the weighted-UniFrac distance matrices, were carried out using the “adonis” and “anosim” function of the “vegan” package in R. Wilcoxon rank-sum test was used to identify the microbial populations with statistically differential abundances among the wellheads and downhole of the water-injection and oil-production wells. Correlations between the relative abundance of bacterial or archaeal OTUs and the shared OTU proportions with permeability of oil-bearing strata were analyzed using the “ggplot2” package in R. Mantel test and ADONIS analysis were used to detect correlations between environmental variables and microbial community compositions using the “vegan” package in R. Based on the long length of the first axis of detrended correspondence analysis (DCA), the unimodal ordination method, canonical correspondence analysis (CCA) was performed to elucidate the relationship between environmental factors and OTU-level microbial communities using the “vegan” package in R.

### Sequences accessibility

The raw reads obtained by Illumina MiSeq sequencing were deposited in the Sequence Read Archive (SRA) at the National Center for Biotechnology Information (https://www.ncbi.nlm.nih.gov/bioproject/PRJNA489604).

## ACKNOWLEDGMENTS

This work was funded by the National Natural Science Foundation of China (Grant No. 31500414 and 41773080), Shandong Provincial Agricultural Fine Species Project (2019LZGC020), National Key Research and Development Project (Grant No. 2018YFA0902101), Jining key research and development project (Grant No. 2019ZDGH019), and National High Technology Research and Development Program of China (Grant No. 2013AA064402). We thank Prof. Ying Zhang, Institute of Applied Ecology, Chinese Academy of Sciences, Shenyang, China, and Dr. Jianlong Xiu, Research Institute of Petroleum Exploration and Development (Langfang), Langfang, China, who provided technical support in core-flooding test. We also thank Fang Liu and Xiaobo Liu, No. 2 Oil Production Plant, PetroChina Daqing Oilfield Limited Company, Daqing, China, for their assistance in sample processing.

